# circuitSNPs: Predicting genetic effects using a Neural Network to model regulatory modules of DNase-seq footprints

**DOI:** 10.1101/337774

**Authors:** Alexander G. Shanku, Anthony Findley, Cynthia Kalita, Heejung Shim, Francesca Luca, Roger Pique-Regi

## Abstract

**Motivation:** Identifying and characterizing the function of non coding regions in the genome, and the genetic variants disrupting gene regulation, is a challenging question in genetics. Through the use of high throughput experimental assays that provide information about the chromatin state within a cell, coupled with modern computational approaches, much progress has been made towards this goal, yet we still lack a comprehensive characterization of the regulatory grammar. We propose a new method that combines sequence and chromatin accessibility information through a neural network framework with the goal of determining and annotating the effect of genetic variants on regulation of chromatin accessibility and gene transcription. Importantly, our new approach can consider multiple combinations of transcription factors binding at the same location when assessing the functional impact of non-coding genetic variation.

**Results:** Our method, `circuitSNPs`, generates predictions describing the functional effect of genetic variants on local chromatin accessibility. Further, we demonstrate that `circuitSNPs` not only performs better than other variant annotation tools, but also retains the causal motifs / transcription factors that drive the predicted regulatory effect.

**Availability:** http://github.com/piquelab/circuitSNPs

## 1 Introduction

Regulation of gene expression is governed by the binding of transcription factors to specific motifs along the genome. These DNA sequences constitute a grammar that is still not yet completely understood (Weingarten-Gabbay and Segal, 2014). Recent experimental techniques have substantially increased the ability to identify and locate these regulatory regions, catalog the interacting factors and their sequence motifs, and annotate genetic variants that could affect chromatin accessibility and regulation of gene transcription (ENCODE Project Consortium, 2012; Boyle *et al.*, 2012; Neph *et al.*, 2012; Buenrostro *et al.*, 2013).

In addition to experimental approaches, a number of computational studies have been undertaken in recent years. Early studies focused their efforts on cataloging predictions of transcription factors binding locations in yeast, fly, and humans (Elemento and Tavazoie, 2005; Xie *et al.*, 2009, e.g.) using sequence models and sequence conservation across orthologous regulatory regions for different species. As new experimental assays that measure chromatin state and accessibility became available, new computational models were developed to integrate different types of information. One of these examples, and also the starting point for the new method we are proposing, is called CENTIPEDE (Pique-Regi *et al.*, 2011). CENTIPEDE uses both sequence specificity and data from genome-wide chromatin sensitivity assays in an empirical Bayesian approach, ultimately generating a posterior probability that a motif in a regulatory region is actively bound by a factor. In this framework, the prior probability is described using a logistic function of a score derived from a position weight matrix (PWM) model determining the binding strength of the transcription factor, while the likelihood is jointly modeled by the number of DNAse-seq reads observed and their spatial distribution surrounding each motif. Our group further developed the CENTIPEDE framework (Moyerbrailean *et al.*, 2016a) and applied it to the task of examining how transcription factor binding might be affected by genetic variants in these key regulatory regions. In our previous work, we recalibrated the sequence models (PWMs) using DNAse-seq data and orthologous sequences, and identified over 3 million variants predicted to significantly alter the prior odds of binding by ≥ 20 fold; those variants now being described as“centiSNPs”http://genome.grid.wayne.edu/centisnps/.

Other methods using machine learning approaches have been proposed. The DeepBind model (Alipanahi *et al.*, 2015) uses a convolutional neural network to learn de novo sequence motifs from sequence data, training on multiple data types including: PBM, ChIP-seq, and HT-SELEX. In addition to identifying putative regulatory motifs, the authors also used their method to predict detrimental variants, creating “mutation maps” that indicated the predicted binding effect of a given variant. More recently, Lee *et al.*, 2015 proposed deltaSVM, a modification of the gkm-SVM model (Ghandi *et al.*, 2014, 2016). Here, the authors use a support vector machine to learn regulatory regions solely from DNA sequence data. By examining the change in the learned model’s output scores between the reference allele and its variant, the authors generate a prediction on the genetic variant effect, and validate it against experimentally determined DNaseI-sensitive QTLs (dsQTLs). DeepBind and deltaSVM utilize as input DNA sequence, only; while centiSNP predictions use both sequence and footprinting information. The footprint shape can effectively remove false positives by further integrating experimentally derived information in the model. Here we have developed a new method that improves the centiSNP approach by modeling effects of multiple motifs in a region. Our approach combines the output of CENTIPEDE and CentiSNPs predictions to also take into account that the binding potential of some factors and their ability to open chromatin may depend on specific combinations of motifs in a region as in a logic circuit, thus moving computational predictions one step further towards representing the complexity of of chromatin accessibility regulation.

## 2 Approach

To better model the underlying biology of transcription factor binding, and to improve the quality of computational predictions on genetic variant effects, we extended the CENTIPEDE and centiSNPs approaches we developed in Pique-Regi *et al.*, 2011 and Moyerbrailean *et al.*, 2016a, moving from a singular motif prediction to a method that models multiple combinations of motifs and the resultant synergistic binding effects. Here, we propose a novel neural network approach that uses deep learning framework, with the intuition being that many factors work cooperatively and competitively in their regulatory roles. Specifically, we envision a model that treats these combinations of factors like multiple switches and logic gates within a circuit. Our model starts from a compendium of high quality footprints generated across several cell types and tissues Moyerbrailean *et al.* (2016b) and predicts genetic effects. Here we applied it to chromatin accessibility data collected in LCLs, but it can be used to generate predictions for any cell-type and cellular context for which chromatin accessibility data is available. The goal of our method is to learn the configuration of these switches and circuits, and, ultimately in doing so, learn which genetic variants may alter the cis-regulatory circuit specific to a target cell-type or cell-condition.

## 3 Methods

### 3.1 Neural Network Modeling of Chromatin Accessibility

Our aim is identifying genetic variants that affect transcription factor binding in a specific cellular context, and thus causally affect the surrounding chromatin structure that may lead to changes in gene transcription. To achieve this we use a two step approach: We first propose a neural network classifier whose input features are individual transcription factor footprints, and where each input example represents a genomic region. We then use a binary variable at each motif to indicate whether a footprint associated with that motif has been detected in that genomic window. The output or class variable for each window indicates that the chromatin is either open or closed - the exact value of the threshold for considering a region open is not very important as long as it is a reasonable value. The resulting neural network is trained to predict chromatin status using patterns of footprint associated with that location.

In the second step, we use CentiSNPs annotation and the trained network to identify which genetic variants are likely to impact transcription factor binding given the neural network “circuit” context. To this end we use the chromatin status predicted for these genomic regions as the two classes in our model (open or closed chromatin), we turn next toward populating our training data using a scheme that jointly incorporates and encodes both sequence information and high-resolution footprints. We first make use of the footprinting analysis and variant effect predictions conducted in Moyerbrailean *et al.*, 2016a. Under their centiSNPs model, the authors first scanned the human genome using 1,949 PWM sequence models, identifying and recording the locations and sequences of all motif matches. DNase-seq data was extracted from 653 samples (from 153 unique tissue/cell types) obtained from the ENCODE and Roadmap Epigenomics projects (ENCODE Project Consortium, 2012; Roadmap Epigenomics Consortium *et al.*, 2015), and in spatial conjunction with the aforementioned sequence matches, transcription factor binding was predicted using a modified implementation of the CENTIPEDE software package (Pique-Regi *et al.*, 2011; Moyerbrailean *et al.*, 2016a). From their analysis the locations of predicted DNase footprints for each motif were determined by a posterior probability threshold of *P >* 0.99, resulting in 6,993,953 non-overlapping footprints across 1,372 motifs.

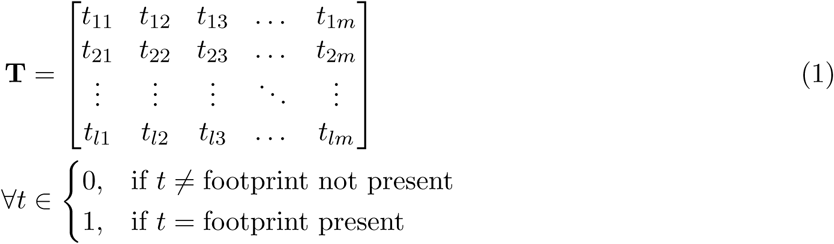

Making use of the locations of these footprints, we construct the training data matrix, **T** = *t*_*lm*_, with *l* indexing a row for each chromatin window and *m* indexing the motif. Each column, or feature, of the training data represents a motif, and each row, or example, corresponds to a genomic location.

To populate the matrix, **T**, we utilize a binary scheme where each element ∈ {0, 1}, where a value at *T*_*l,m*_ = 1 represents a footprint present at location *l* for motif *m* (See Equation (1)). Similarly, *T*_*l,m*_ = 0 when motif *m* does not have a corresponding footprint at location *l*. The vector of outputs, ŷ, can take binary labels, class labels, or continuously valued labels. Visually, this is depicted in Figure 1.

**Figure 1:**
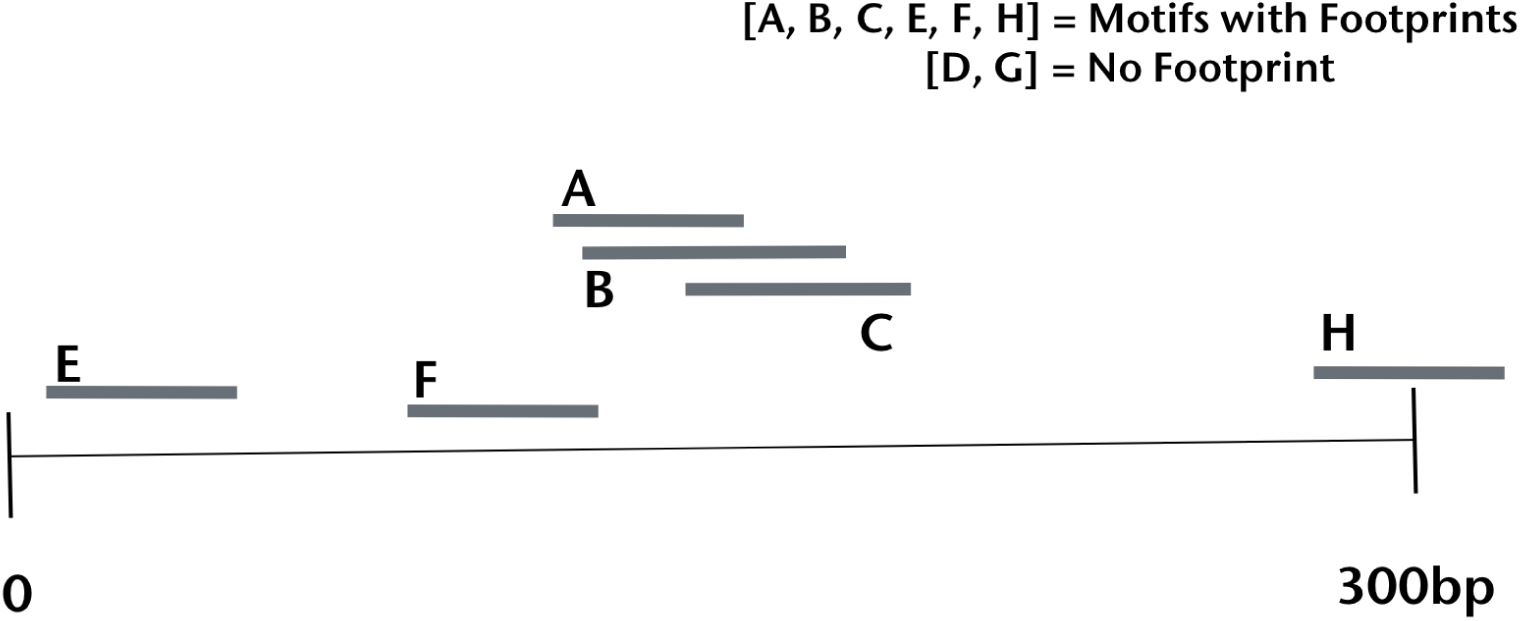
In this simplified visual representation of our model’s training data, we use previously annotated footprinting data to determine which motifs have footprints in a specified region. Footprints are represented as horizontal lines labeled **A** though **H**. Data are encoded in a binary format, such that motifs with verified footprints are labeled a “1”; conversely if a motif does not have a representative footprint, that element in the “Footprint vector” is labeled a “0”. *E.g.* in this figure we depict annotations for eight motifs, {**A**, **B**, **C**, *…,* **H**}, thus the resultant encoding would be *V*_*foot*_ = [1, 1, 1, 0, 1, 1, 0, 1], as footprints of motifs **D**and **G** are not present in this window.

Each row vector of *T* is thus an input to our neural network and its class label is either 0 or 1, depending on the state of chromatin accessibility at that location, closed or open, respectively. In training, data are sampled, without replacement, such that 75% are used in model training, 12.5% in model validation, and 12.5% as test data to determine the trained model’s performance.

#### 3.1.1 Neural network architecture

To model the relationship between open chromatin and footprints we use a neural network approach, which is well suited to learn an arbitrary logic function between the combinations of observed footprints present in a genomic window and their potential to affect that region’s chromatin accessibility. To this end, our neural network consists of three layers: two fully connected hidden layers and a single output layer. To explore a range of hidden units in the hidden layers, we train ten replicates of each model using the following combinations of hidden layer units: (250,40), (50,10), (10,5), (5,3), (1,0), and (0,0). The final layer in the network maps the output to a value ∈ (0, 1), essentially a probability describing the chromatin state. The total number of trainable parameters are dependent on the number of hidden units (See Section 4.1 and table S1)

Each hidden layer uses a ReLU activation function, while the output layer uses a sigmoid activation function. All models used binary-crossentropy as a loss function and the adaptive learning rate Adadelta optimizer (Zeiler, 2012). We allow our model to train for 50 epochs, while monitoring the validation loss and stopping the training if loss does not decrease for 3 consecutive epochs. Development was accomplished using the Keras library and Theano as the backend. (Chollet *et al.*, 2015; Al-Rfou *et al.*, 2016).

### 3.2 circuitSNPs: Integrating genetic variant effects

Once we learn the function that connects footprint combinations with open chromatin in a specific cell-type, the goal is to predict the effect of genetic variants on the overall output. We denote these genetic variants as `circuitSNPs`. To achieve this we first consider the effect of a genetic variant on the footprint inputs using the computational predictions generated by Moyerbrailean *et al.*, 2016a. In addition to the footprint annotations used in creating our training data (see Section 3.1), the authors have also annotated 8,011,245 SNPs from the 1000 Genomes Project (1000 Genomes Project Consortium *et al.*, 2012). These annotations consider if the genetic variant is in a TF footprint and if it is predicted to alter the prior odds of binding ≥ 20-fold (centiSNP). The annotation is directional, thus predicting whether the reference allele increases or decreases TF binding at that location. Of 5,810,227 variants in footprints, 3,831,862 were predicted to significantly alter TF binding.

Using the centiSNPs annotation, we construct the core of the `circuitSNPs` model, by taking into consideration the direction of predicted effect on TF binding and the model we learned in the previous section. Specifically, we ask whether binding increases or decreases for a given transcription factor when the reference/alternate allele for a genetic variant is present in the TF motif. With this in mind, the `circuitSNPs` model generates a prediction that is obtained by calculating the log-odds difference between a “reference prediction” and an “alternate prediction” while taking into consideration its specific regulatory context considering all the footprints in the region when evaluating the neural network output. More specifically, for each SNP we generate a prediction for the “reference” and “alternate” matrix as follows: 1) Similar to how we populated our training data in Section 3.1, we again create a matrix, **X** where, at each SNP’s location, we determine all motifs with an overlapping footprint and set that matrix element *X*_*l,m*_ = 1. 2) For the reference prediction, we “turn off” footprints if the reference allele significantly decreases binding. That is, elements in each row vector equal to 1 are set to 0 (See Figure 2). 3) Similarly, for the alternate prediction, we set elements that the alternate allele significantly decreases binding to a zero. 4) We then use our trained neural network and classify bot the reference and alternate vector to generate an output. 5) Finally, to generate the `circuitSNPs` prediction, we calculate the log-odds difference between the reference and alternate prediction.

**Figure 2:**
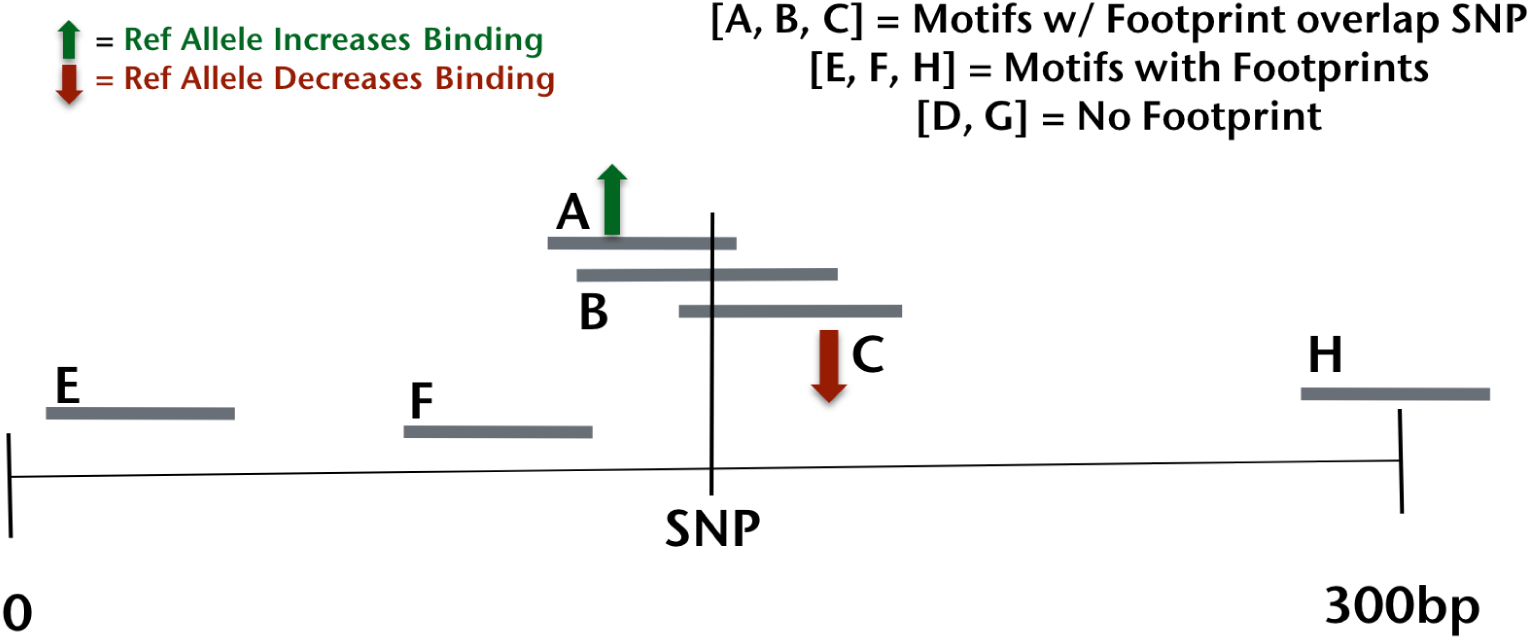
To generate the `circuitSNPs` predicted effects for each genetic variant we only consider footprints that overlap with that variant or are within very close proximity and can interact with each other. We summarize this information as a vector *V*_*F*_ with the same dimensions as that one used for our trained neural network. We then modify this vector, creating two new vectors, *V*_*R*_ and *V*_*A*_, such that each element in *V*_*R*_ that corresponds to a variant in that motif decreasing binding for a reference allele is set to 0, and each element in *V*_*A*_ that corresponds to a decrease in binding for the alternate allele is set to 0, leaving all other elements unchanged. In this figure, *V*_*F*_ = [1, 1, 1, 0, 0, 0, 0, 0], becomes *V*_*ref*_ = [1, 1, 0, 0, 0, 0, 0, 0] and *V*_*alt*_ = [0, 1, 1, 0, 0, 0, 0, 0], corresponding to the predicted effects of the variant on motif **A** and **C**.

### 3.3 circuitSNPs prediction validation

We trained and validated our model on the regions of open and closed chromatin described in Lee *et al.*, 2015, and validated its ability to predict a variant’s effect on binding against two data sets containing variants associated with inter-individual variation in chromatin accessibility. The first validation set is a subset of the Degner *et al.*, 2012a dsQTLs, as described in (Lee *et al.*, 2015). For our second validation, we used 8,590 dsQTLs identified using RASQUAL (Kumasaka *et al.*, 2016) on the data from Degner *et al.*, 2012a. In the first test, following the procedure described in Section 3.2, we generate predictions on the dsQTLs that are used for evaluation in Lee *et al.*, 2015. This dataset contains a small set of all dsQTLs (579 out of 8,902 from Degner *et al.*, 2012a) and 27,735 additional negative controls. Our second validation approach examined the correlation between the associated RASQUAL effect-size scores of these data (*π*-score) and our circuitSNP predictions.

### 3.4 Using circuitSNP predictions and the retained motifs to examine co-binding events in DNAse-sensitive QTLs

Conditioning on whether or not the variant is in a true validated dsQTL, and by filtering circuitSNP predictions as those ≥ *|*3.0*|*, we construct four count matrices and use Fisher’s exact test at 10% FDR to determine those motifs that are enriched for co-occurrences with other motifs.Specifically, a motif-motif co-occurrence is described by both motifs being present in a given genomic window. We construct a 2×2 table for each motif, describe the co-occurrences with all other motifs, with each cell being populated using the following criteria: 1) the motif is active at a variant with a significant circuitSNP prediction and that variant is present in the experimentally validated dsQTL, 2) the motif is active at a variant with a non-significant circuitSNP prediction and that variant is present in the experimentally validated dsQTL, 3) the motif is active at a variant with a significant circuitSNP prediction and that variant is not present in the experimentally validated dsQTL, and 4) the motif is active at a variant with a non-significant circuitSNP prediction and that variant is not present in the experimentally validated dsQTL.

## 4 Results

### 4.1 Neural Network Model Training and Validation

The first part of the our proposed approach is to learn the regulatory models that are composed of combinations of footprints. In this training phase, we utilize a set of 134,304 training DNase-I hypersensitivity regions defined in (Ghandi *et al.*, 2014; Lee *et al.*, 2015), with 22,384 positive example chromatin-accessible regions of size 300bp, and an additional 111,920 GC-matched negative sequences of the same size with a closed chromatin profile. Alternatively, for a new tissue we can similarly construct a set of regions in a similar way as defined by the ENCODE project. To determine the ability of our trained model to predict genomic regions of open chromatin, we examined the metrics of the precision recall curve and area under the precision recall curve (auPRC). An example of one training replicate is shown in Figure S1. Based on the ability of circuitSNPs to integrate the signal of a footprint’s presence across hundreds of motifs, we note that footprinting data, alone, allows us to predict the chromatin state in that region with an auPRC of 0.74 and 98% precision at 10% recall. While learning open chromatin regions is part of the process of building the neural network the final objective of our method is to predict genetic variants that affect chromatin regulation.

Using the model parameters and training procedure described in (Section 3.1.1 and table S1), we observed the effect of these parameter combinations on auPRC and precision at 10% recall on our predictions when applied to the dsQTL data set described in Lee *et al.*, 2015 (Ghandi *et al.*, 2014, 2016) (See Section 3.3 and Degner *et al.*, 2012a). In Figure 3 we show the mean values of the ten replicates for each parameter combination, along with the associated standard error. For the six replicates, the respective mean auPRC and standard error is (0.149, 0.003), (0.182, 0.002), (0.196, 0.004), (0.204, 0.001), (0.185, 0.004), (0.180, 0.001). The precision @ 10% recall for these replicates is (0.379, 0.023), (0.557, 0.011), (0.657, 0.011), (0.687, 0.009), (0.639, 0.011), (0.605, 0.004), respectively. For comparative purposes, the gkmSVM metrics for this data set is plotted as well (auPRC = 0.193, prec. at 10% recall = 0.538) (Lee *et al.*, 2015). We find that our model’s performance decreases as we increase the number of hidden units in the hidden layers. For example, under a model with 200 units in the first layer, and 40 units in layer #2, we find an auPRC and precision at 10% recall of ∼ 0.15 and 0.375, respectively, whereas the same metrics under the “5*|*3” model are ∼ 0.2 and 0.7, respectively.

**Figure 3:**
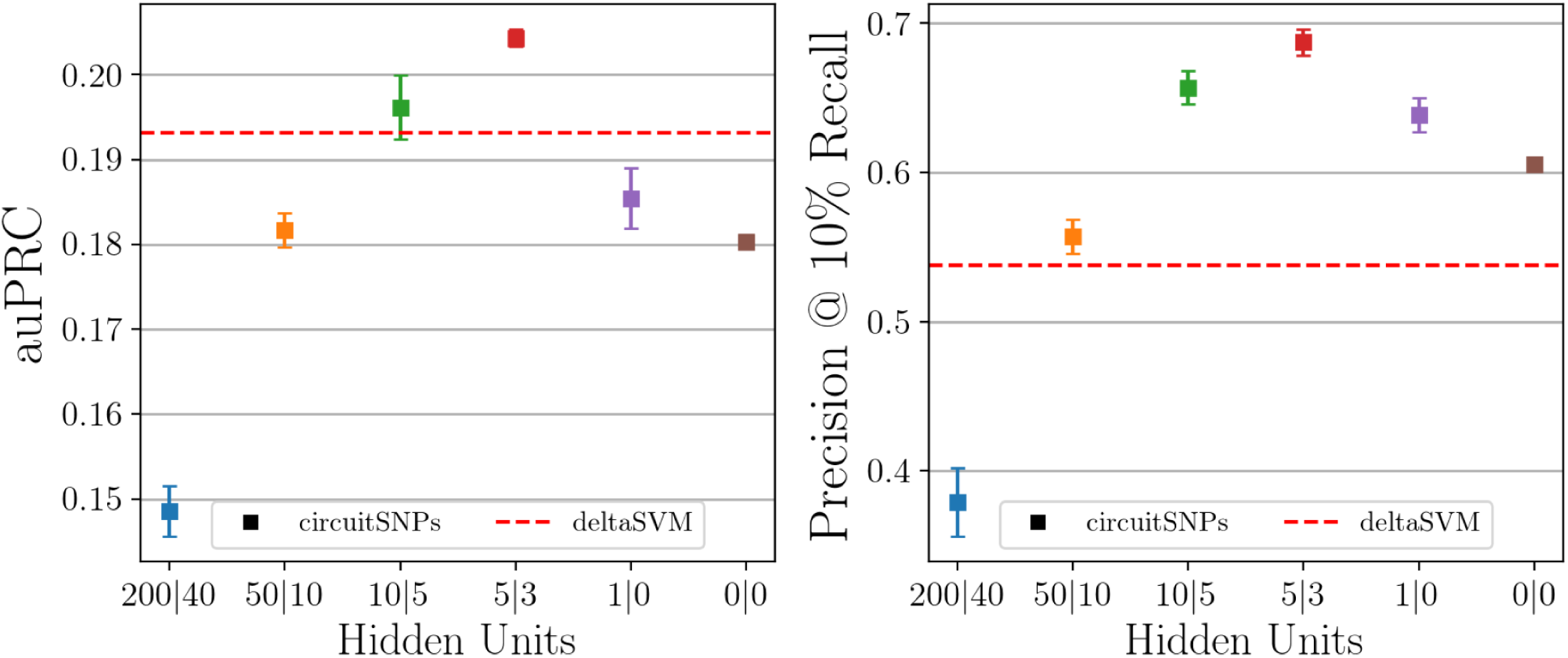
To investigate how the number of hidden layer units affects our model’s performance, we repeated our cross-validation procedure 10 times for each of the six hidden layer parameter combinations. The left panel shows the area under the precision recall curve (auPRC). The right panel shows the precision (or positive predictive value) achieved at 10% recall. Error bars are calculated standard error and dashed horizontal red lines show gkm-SVM scores for both metrics as reported by the authors.

Next, we examined how our circuitSNP predictions correlated with RASQUAL *π* statistics for 519,387 sites identified as dsQTLs. Using an FDR of 10% for *π*, we retained 8590 variants to analyze. Using the ten replicate models for each of the six hidden unit parameter combinations, we calculated the correlation coefficient (*r*) means and standard error (Figure 4 left). From the greatest to least complex parameterization, the mean *r* and SE are (0.2409, 0.0034), (0.2736, 0.0016), (0.2866, 0.0035), (0.2937, 0.0013), (0.2857, 0.0014), (0.2836, 0.0003), respectively. Again, we note that the “5*|*3” model shows the greatest mean *r* value of the model replicates tested. Using the best model parameterization, we examined how well our large-effect predictions (≥ *|*3.0*|*) agreed with the RASQUAL allelic effect direction (94.8% agreement) and show a stronger correlation (*r* = 0.81, p≤ 4.3 × 10^−56^), (Figure 4 right) for those events where the model predicts a stronger effect.

**Figure 4:**
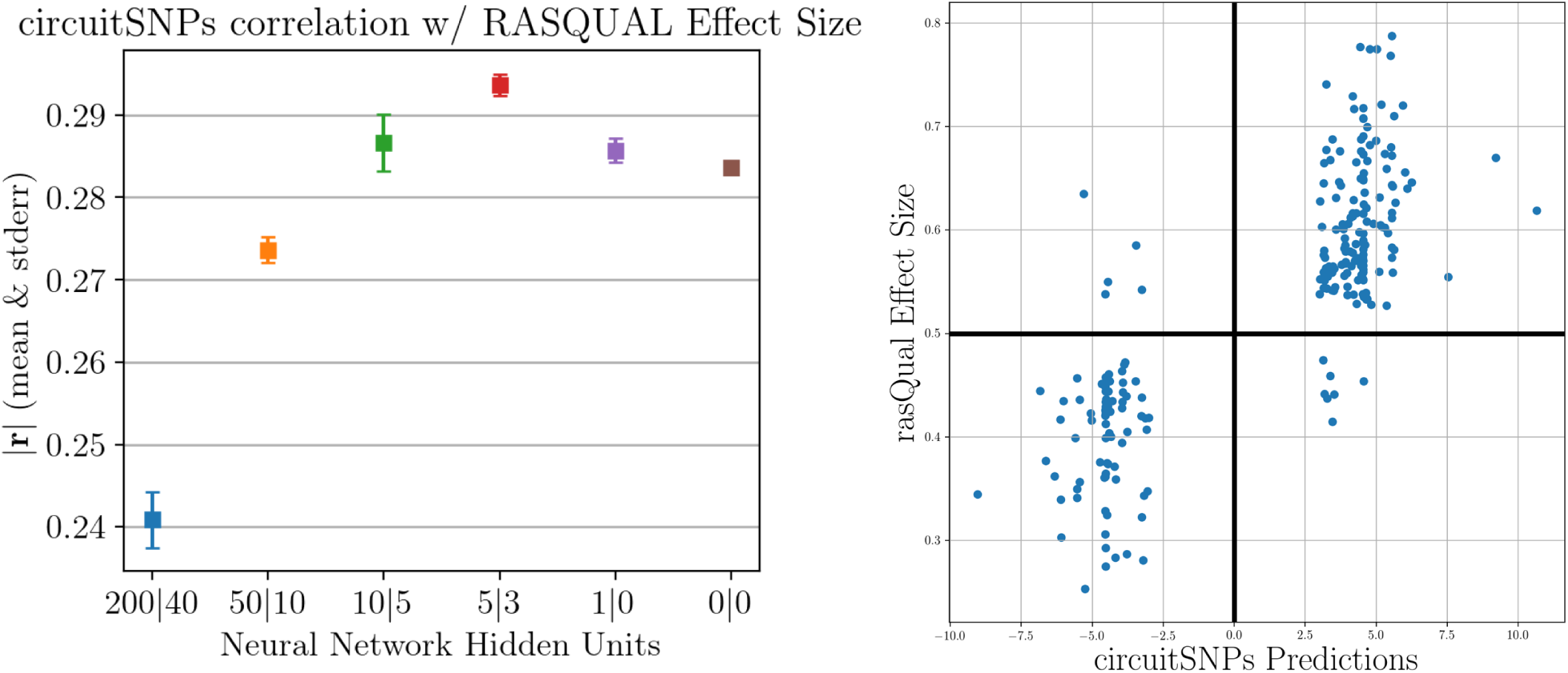
Using the replicate trainings we plot the circuitSNPs predictions correlations to the RASQUAL effect sizes (Degner *et al.*, 2012b; Kumasaka *et al.*, 2016). On the left panel: for each parameter combinations, an error plot shows the mean Spearman’s correlation *r* value for 10 model replicates at that corresponding parameterization. Each replicate consists of 150,000 circuitSNP predictions and RASQUAL score pairs. On the right panel: Scatter plot of circuitSNPs predictions and a set of experimentally derived effect sizes from Degner *et al.*, 2012b; Kumasaka *et al.*, 2016. Shown here are circuitSNP predictions and RASQUAL score pairs, filtered for small effect size, that yield a highly significant *r* = 0.82 (*p* ≤ 4.3×10^−56^)

We also examined the effect of increasing the spatial signal of neighboring footprint signals when creating the “reference” and “alternate” vectors prior to making a circuitSNPs prediction (See Section 3.2). We used the “5*|*3” model and tested four window sizes around the variant where a footprint must be present in order to be included in the model input: 300bp, 100bp, 10bp, and 0bp (meaning the footprint must overlap the focal variant). Interestingly, we show that the best results are obtained by using a model where we condition only on those motifs whose footprints overlap the variant. In Figure S4 we show the means and standard error of the auPRC and precision at 10% recall for the four windows investigated.

### 4.2 Generalizability of the model

The circuitSNPs model presented here is trained on chromatin footprints from derived from multiple tissues. In this sense, our model is very generalized in training, but may be applied to tissuespecific tasks. We demonstrate this specifically in Section 4.1, where we validate our model against lymphoblastoid cell line data. In some instances, it may be advantageous to train and utilize the circuitSNP model in a single tissue, or in a selected set of tissues chosen *a priori*. However, in the case we observe here, where the data are dsQTLs and task is predicting the genetic variant effects in LCLs, we note that training across many tissues and multiple experiments generates better overall performance. Table S2 demonstrates the performance difference between three implementations of the circuitSNPs model when employed to predict variants in LCLs: 1) a model trained and applied using only data from 18 LCL tissues 2) a model trained using all 153 tissues, but applied as inputs populated by 18 LCL tissues, and 3) a model trained and applied using 153 tissues. This results show that we can leverage more footprint information by combining the entire compendium of footprints as it will have less false negatives and that the neural network approach effectively can learn which are relevant specifically for LCLs.

### 4.3 Variant Directionality

When developing circuitSNPs, we hypothesized that the ability to integrate the combinatorial effects from a large number of motifs bound in a genomic window is a better model to analyze variant effects, versus looking at predictions generated from individual motif models. To test this hypothesis, we use the analysis in Section 3.3 and examine the agreement in the direction of the allelic effect between the circuitSNPs predictions and the RASQUAL scores, compared to the individual motifs-derived predictions and RASQUAL scores. We obtain first the number of circuitSNP predictions that agree with RASQUAL scores at each variant predicted, as a fraction of all circuitSNP/RASQUAL predictions. Then, for each motif represented in our model, we similarly calculate the fraction of a motifs predicted effect that agrees with the RASQUAL score, versus the total number of times that a motif/RASQUAL prediction appears in our data. Using this metric, we find that circuitSNPs ability to incorporate and utilize the sequence and footprint information from all motifs provided for greater agreement versus the individual motifs, alone. This is particularly evident for variants in footprints for c-Ets-1(p54), Ehf, Elf4, Elk3, ERG, Etv1, MAFA, and PDEF (represented by points falling below the (0,1) line in Figure 5).

**Figure 5:**
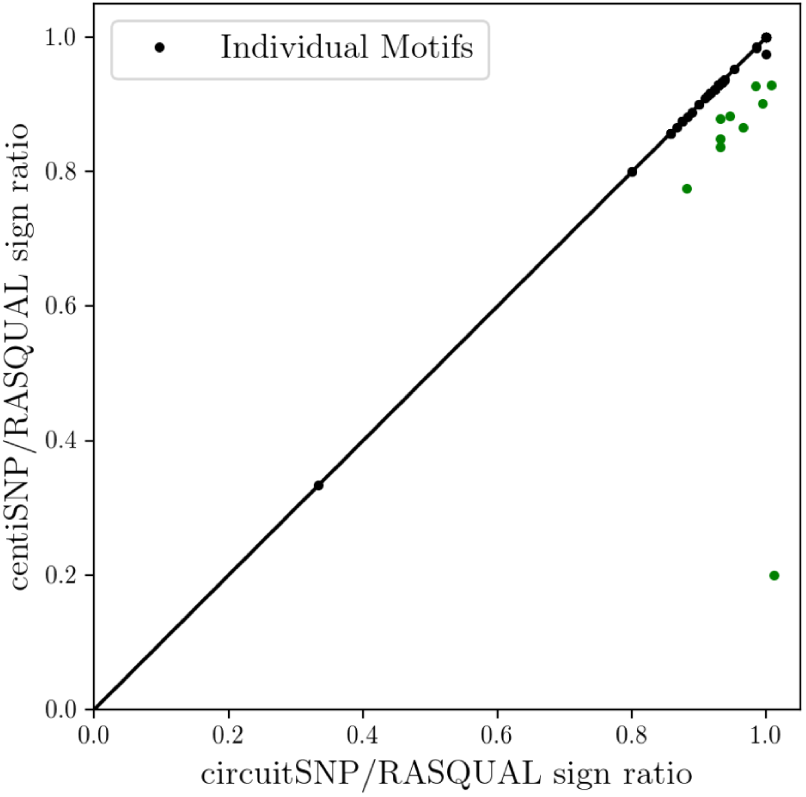
The circuitSNPs prediction, which integrates footprints and sequence information from all motifs at the variant’s location, better captures the directional variant effects than using individual motif predictions, alone. The green dots in the scatterplot show the plotted ratios of circuitSNP sign direction to RASQUAL sign direction, vs. individual motif prediction direction to RASQUAL sign direction. Points falling below the zero-one line represent cases where the circuitSNP prediction performs better than the motif alone.

### 4.4 Motif Co-binding Enrichment

One of the goals of our model is that we retain the causal motifs at each variant and prediction to facilitate downstream analysis and a better understanding of which motif combinations may be more impacted by genetic variation. Using the data set presented by Lee *et al.*, 2015, we sought to identify significantly enriched co-binding partnerships between motifs associated with allelic effects observed in dsQTLs. Using the criteria specified in Section 3.4, we tabulated 7,529,536 co-occurrence counts (4 cells ×1372^2^ motif co-occurrences) to compare the co-occurrences of motifs in our circuitSNP predictions to the co-occurrences observed in the experimentally derived dsQTL dataset. We determined significant motif pairings in the dsQTL set using Fishers exact test at 10% FDR (Benjamini Hochberg). We found that 37 motifs pairings, representing 21 unique motifs, were either depleted or enriched for their respective co-occurring partners (14 unique motifs), see Figure 6A. We then tabulated matrices of co-occurrences as described in Section 3.4 for both our significant and non-significant circuitSNP predictions and calculated the log10 odds ratios to see if our neural network can also capture these enrichments when we predict the impact of genetic variation. We found that our circuitSNPs predictions agreed with the validated dsQTL enrichments in 21 of the 37 (56.75%) co-occurrences Figure 6B. We note, however, that of the 15 pairs that are not in agreement between the dsQTLs and our predictions, 12 of these 15 (80%) occur in a single motif, MA0028.1. Excluding this motif, our circuitSNPs predictions are in agreement with 17 of the 20 remaining co-occurrence events (85%). In contrast, examining dsQTL co-binding against CentiSNP predictions that do not model footprint combined effects we find agreement in only 4 of the 37 (10.8%) pairings Figure 6C.

**Figure 6:**
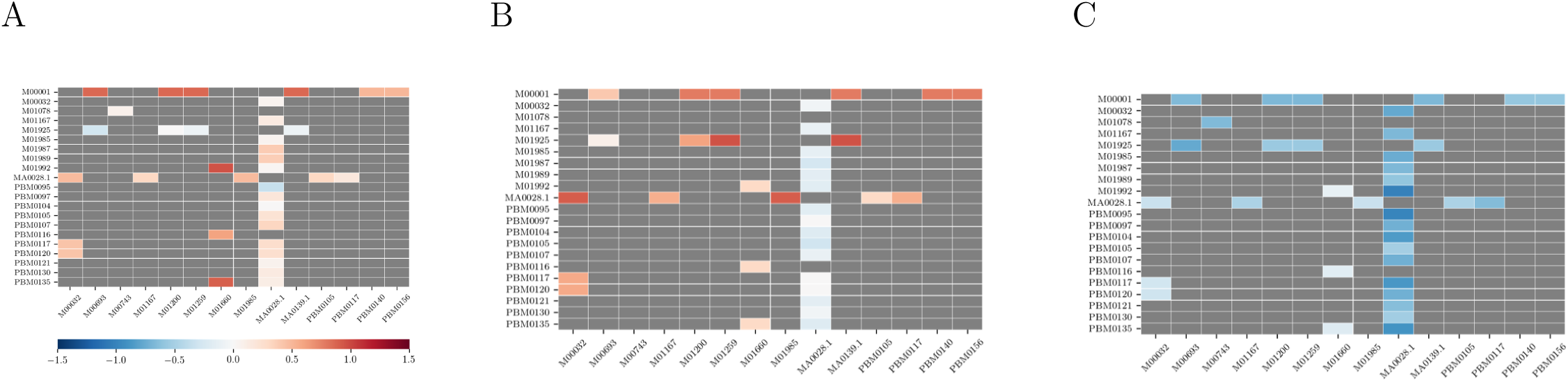
Heat map of co-occurring motif enrichment, colored by the *log*_10_ fold change (A) Co-occurring motifs in the dsQTL data. Of the possible 1,881,012 co-occurrence events between motifs present in the dsQTL data, we found 37 such events where a motif is either significantly enriched or depleted for their respective binding partner. (B) Co-occurring motifs based on our new circuitSNPs predictions recapitulate the enrichments observed on the empirical dsQTL data in panel A. (C) Co-currence of motifs with predicted effects based on CentiSNPs not taking into consideration combinations of motifs in the prediction model fails to recapitulate the enrichments observed in the empirical data observed in A

## 5 Discussion and conclusion

Allelic variation can have profound effects on gene regulation, gene expression and ultimately lead to disease relevant phenotypic consequences(Degner *et al.*, 2012a; Neph *et al.*, 2012; Lee *et al.*, 2015; Alipanahi *et al.*, 2015; Moyerbrailean *et al.*, 2016a). In an effort to determine SNPs causal in regulatory variation, we have developed a hybrid method that predicts the effect of genetic variants in regulatory regions. As many transcription factors may bind in a localized, non-coding region, developing methods that look to exploit and decode this regulatory grammar is increasingly important.

We have demonstrated that by integrating the individual effect a SNP has on TF binding for all motifs present at a genomic location, our method outperforms single-motif based predictions for at least ten motifs active in regions where a variant is shown to affect chromatin state. Our results confirm a more complex binding relationship amongst transcription factors in regulatory regions, and, by modeling that complexity, circuitSNPs may provide more accurate predictions that better capture biologically relevant information.

Predicting the effect of a genetic variant in a regulatory region is still a difficult task in genetics. However, using the circuitSNPs model presented in this study, we have demonstrated that our computationally derived predictions that model not only the underlying effects of many motifs at once, but also simultaneously incorporate their chromatin profile, can outperform contemporary methods that are limited to sequence-based modeling. By using individual genetic effects for each motif as inputs into our neural network, we demonstrate that we can directly retain the motif identities without relying on alternative and complex methods to infer causal motifs. As a result, we have shown that for factors that interact with each other, the effect of genetic variants in their motif sequences, are more readily detectable in our new framework, versus predicting those effects at the level of the individual motif. Finally, we show that unique and complex,co-occurrence events between motif pairs are found in dsQTLs, and that this signal can be accurately predicted and detected using the circuitSNPs model.

## Acknowledgements

We thank Natsuhiko Kumasaka for access to the RASQUAL-based dsQTL analysis used throughout this manuscript and members of the Luca/Pique-Regi group for helpful comments.

## Supplementary information

**Table S1:** circuitSNPs Training Parameterization. (See Section 3.1.1).

| Model | Layer 1<br>Hidden Units | Layer 2<br>Hidden Units | Layer 1<br>Parameters | Layer 2<br>Parameters | Output Layer<br>Parameters | Total Parameters |
| --- | --- | --- | --- | --- | --- | --- |
| 1 | 200 | 40 | 274,600 | 8040 | 41 | 282,681 |
| 2 | 50 | 10 | 68,650 | 510 | 11 | 69,171 |
| 3 | 10 | 5 | 13,730 | 55 | 6 | 13,791 |
| 4 | 5 | 3 | 6865 | 18 | 4 | 6887 |
| 5 | 1 | 0 | 1373 | 0 | 2 | 1375 |
| 6 | 0 | 0 | 0 | 0 | 1 | 1373 |

**Table S2:** Using specific tissues to train and test circuitSNPs. “Train” refers to the origin of the fooptrints used for learning the neural network, and “Predict” to the set of fooptrints and CentiSNPs annotation used to generate the predictions

| Input footprints<br>(Train / Predict) | Area Under<br>Precision Recall Curve | Precision @<br>10 % Recall | Recall @<br>50 % Precision |
| --- | --- | --- | --- |
| LCL / LCL | 0.139 | 0.356 | 0.048 |
| All Tissues / LCL | 0.199 | 0.561 | 0.142 |
| All Tissues / All Tissues | 0.241 | 0.628 | 0.175 |

**Figure S1:**
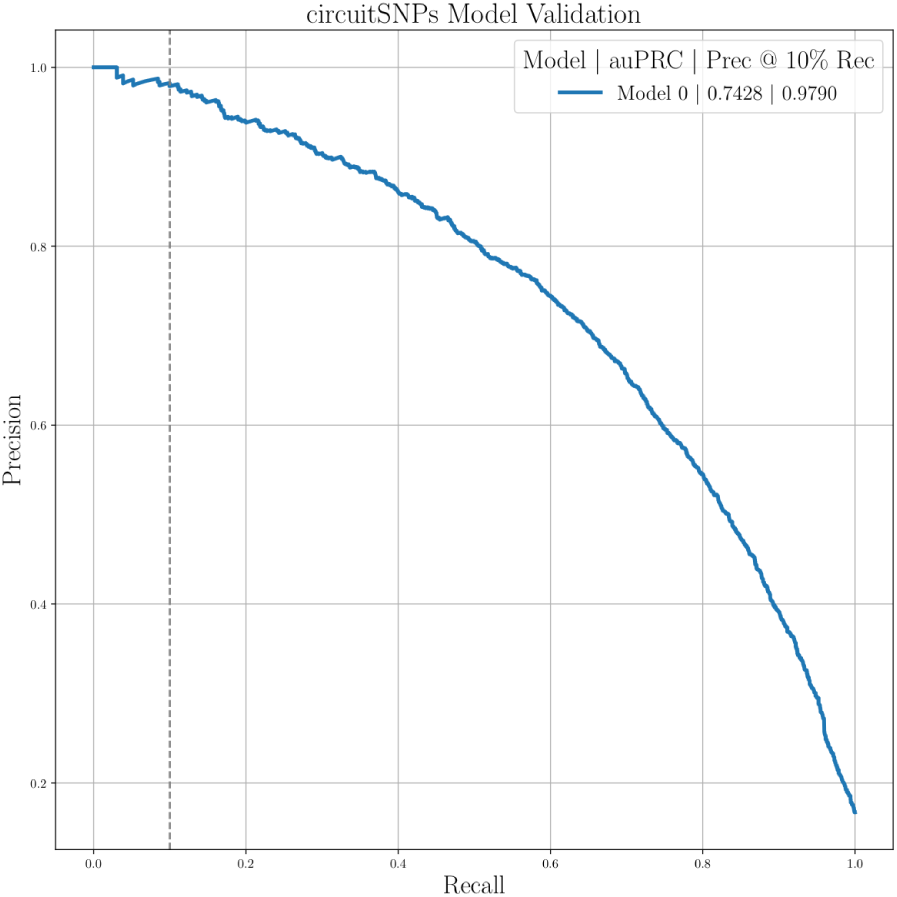
Training the circuitSNP model on footprinting information, alone, allows for accurate prediction of DNAse sensitivity in a genomic window. In one replicate of the training phase, we show an auPRC ofz 0.74 and a precision of 0.98 at 10% recall.

**Figure S2:**
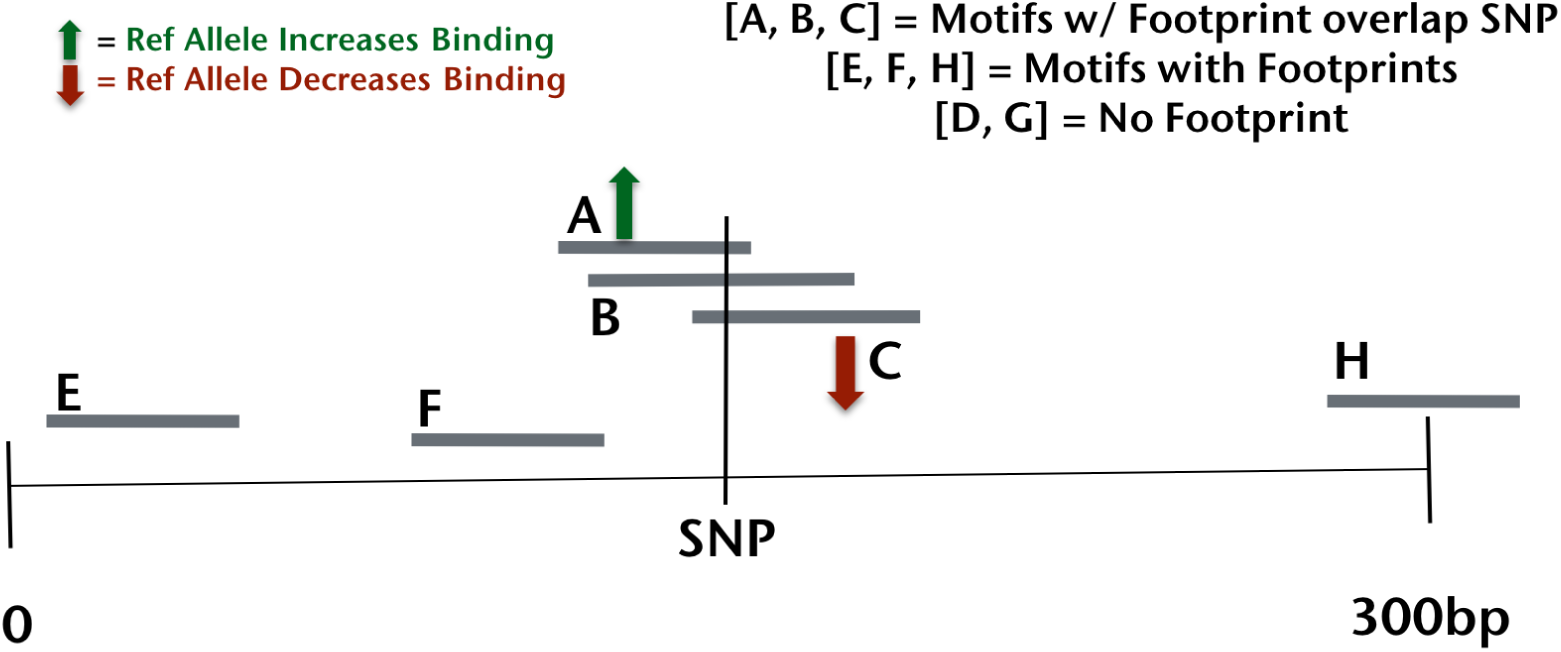
circuitSNPs window model. Here we relax the requirement that a footprint must overlap the focal SNP, as in Figure 2 and section 3.2. We now track all motifs with footprints in the window, the distinction between this approach and the model described in Figure 2 and section 3.2, is that all motifs present in the window are used to create the “Footprint vector”, however, only those that positionally overlap the SNP are manipulated to create the *V*_*ref*_ and *V*_*alt*_ vectors (the same as the non-windowed model). In this figure, *V*_*foot*_ = [1, 1, 1, 0, 1, 1, 0, 1], becomes *V*_*ref*_ = [1, 1, 0, 0, 1, 1, 0, 1] and *V*_*alt*_ = [0, 1, 1, 0, 1, 1, 0, 1], corresponding to the predicted effects of the variant on motif **A** and **C**.

**Figure S3:**
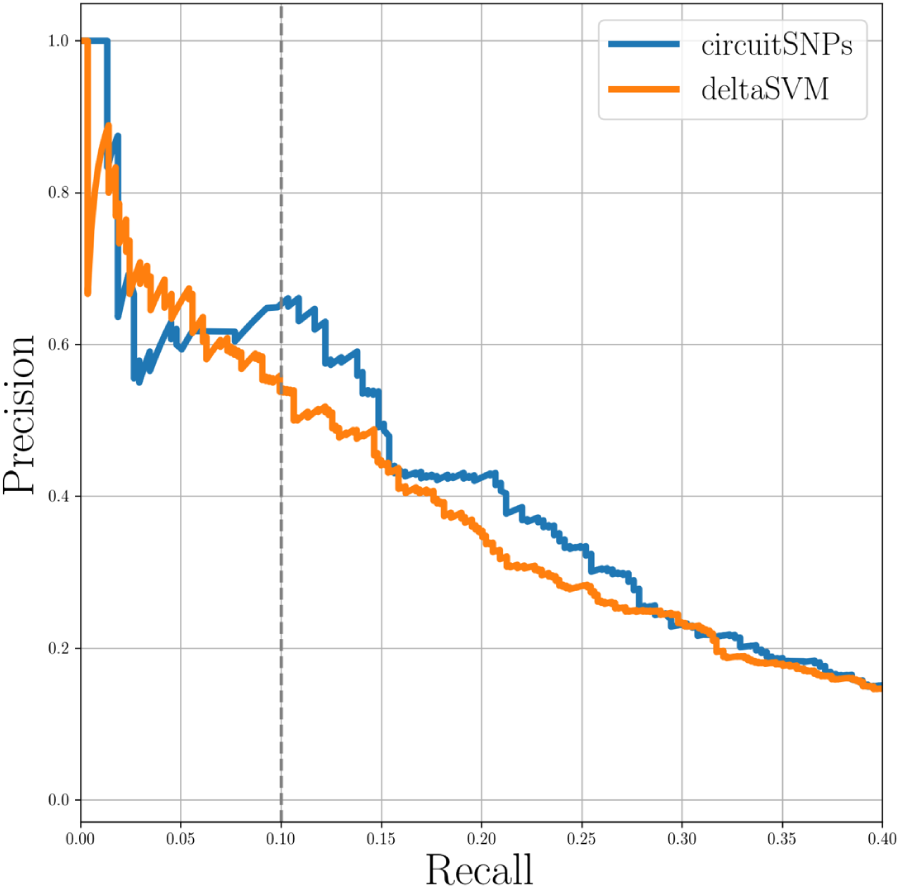
Precision Recall curves comparing circuitSNPs predictions and gkmSVM scores on the dsQTL dataset (Ghandi *et al.*, 2014). We plot the precision recall curve for both the circuitSNPs and gkmSVM models.

**Figure S4:**
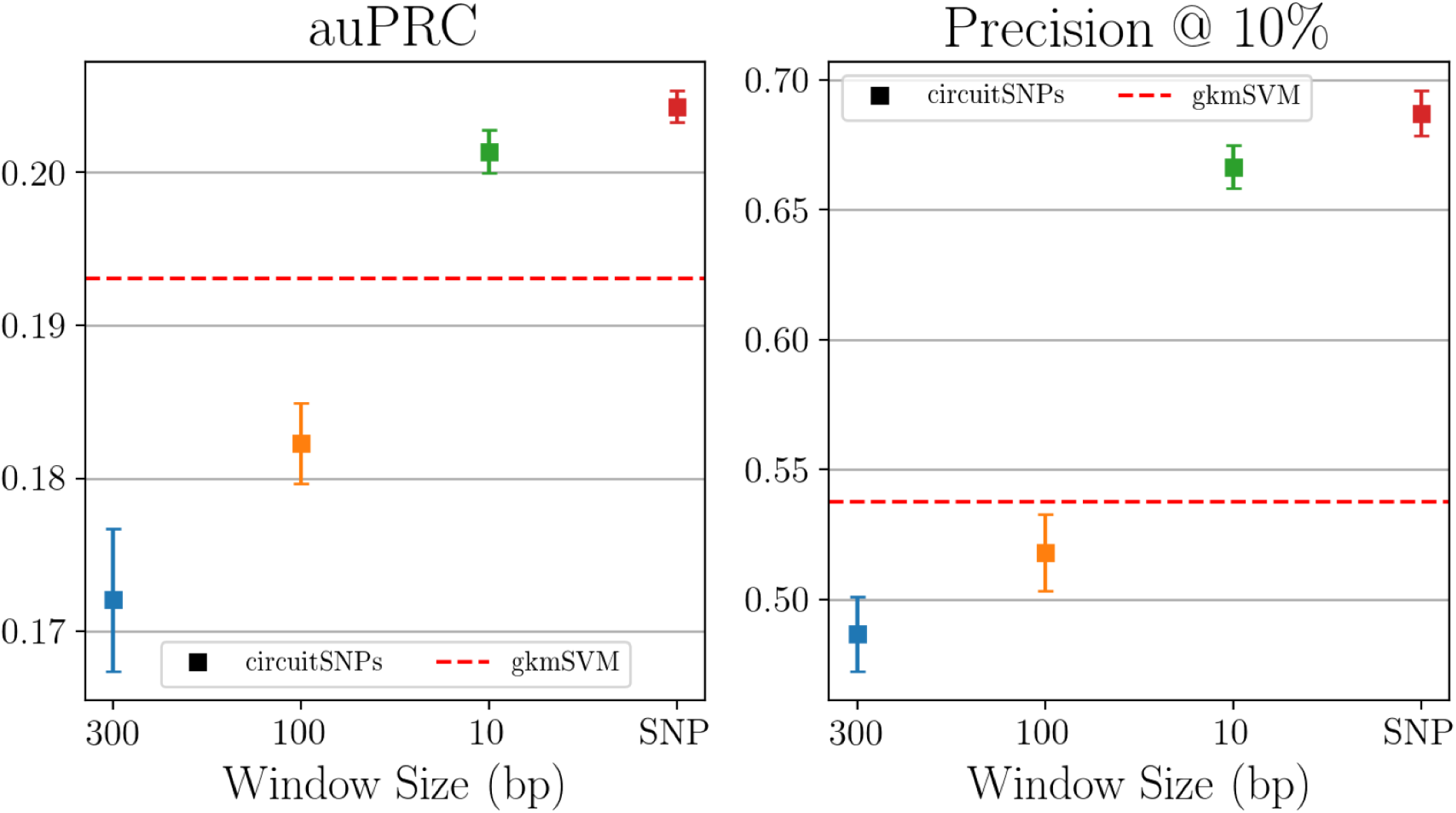
Using the model parameterization that we demonstrated to preform best (Figure 3 and section 3.1.1), we investigated the effect of increased flanking regions around the focal SNP when determining the footprints present. We show that there is a performance relationship associated with the size of the genomic window, with larger window sizes correlated with decreasing performance. Panel “A”: area under the precision recall curve (auPRC), Panel “B”: precision at 10% recall. Error bars are calculated standard error and dashed horizontal red lines show gkm-SVM scores for both metrics.

